# A barcoding approach to phylogenetic classification of Aedini mosquitoes (*Aedes, Ochlerotatus*)

**DOI:** 10.1101/285825

**Authors:** H. Glass, E. Carroll, D. Curley, H. Kienzle, D. A. Yee, S. M. Vamosi

## Abstract

Traditionally, entomologists have used morphological characteristics for mosquito taxonomy and systematics. However, this approach does not take into consideration the genetic relatedness of species. In 2000, the *Aedes* genus of mosquitoes in the tribe *Aedini* was split into two genera (*Aedes* and *Ochlerotatus*), thereby elevating *Ochlerotatus* from subgenus to genus rank, strictly based on morphology of adults. Herein, we use the genetic barcoding marker COI to generate a phylogeny of 65 species of *Aedes, Ochlerotatus*, and *Anopheles* outgroup from almost 900 sequences downloaded from BOLD systems. Our results reveal evidence of non-random, but polyphyletic clustering of *Aedes* and *Ochlerotatus* species, with a monophyletic outgroup. We do find support for the validity of *Ochlerotatus* as an evolutionary unit, although we find insufficient evidence to support its retention as a genus. We suggest that mosquito phylogenetic analyses incorporate a greater number of genetic markers to help clarify our understanding of *Aedini* species classifications, but caution that recent assessments based solely on morphology may be insufficient.

## Introduction

Insects present significant challenges to systematists for several reasons, including their incredible diversity (Labandeira and Sepkoski 1993, Whitfield and Kjer 2008), relatively old age (Dunlop 2010, Whalley 1986), significant variations in diversification rates through time (Barraclough and Volger 2002, Wiegmann et al. 2011), and morphological similarity among congeners, especially in larval specimens (Schultz and Meier 1995). Despite a long history of study as potential vectors of human and animal disease, mosquitoes are no exception to these systematic challenges. There are 3,552 species of Culicidae (Harbach 2016), with the oldest fossil being dated to 90-100 million years old (Borkent and Grimaldi 2004). Although there are few time-calibrated estimates of diversification rates through time, there also is evidence for rapid radiations early in the history of the group (Reidenbach et al. 2009). Adding to these issues of classification, reliable identifications are difficult in various species complexes, especially among some medically relevant species, with some keys requiring male specimens in some groups and female specimens in others (Chan et al. 2014).

Within the Culicidae, taxonomic and systematics relationships are particularly unresolved within the Tribe Aedini. Here, we focus on controversies surrounding the existence of, and relationships between, the genera *Aedes* (Meigen 1818, as cited in Harbach 2016) and *Ochlerotatus* (Meigen 1818, as cited in Harbach 2016). Through a series of morphology-informed phylogenetic studies, Reinert et al. (2000, 2004, 2006, 2008, 2009) elevated *Ochlerotatus* to genus status. This decision was based on the analysis of morphological characteristics of 119 Aedini species across all life stages. The resulting morphology-based phylogeny proposed this change to the previous classification that was based solely on adult mosquito morphology. These revisions to the genera created instant controversy among researchers and led to many journals that focus on these medically important species to suggest caution with adopting the new designations (Reisen 2016). Because many species within the genus *Aedes* are of significant medical importance (e.g., *Aedes aegypti*), redesignation of any species would pose challenges for public health officials in relation to using long standing species names when communicating with the public. In addition, as it has been nearly two decades since Reinert first proposed elevating *Ochlerotatus* to a genus (Reinert 2000), and using a molecular approach to resolving the phylogenetic relationships among these species is long overdue. Indeed, in an editorial about Aedini mosquitoes, Reisen (2016) noted: “As more mosquito sequencing data become available … genetic analyses should be done to confirm these phenotypic groupings.”

DNA barcoding has been promoted as a universal tool for reliable species identifications (Hebert et al. 2003, Hebert et al. 2004), and also as a tool for helping to resolve phylogenetic relationships among species (Hajibabaei et al. 2007, Erpenbeck et al. 2007). The 648 base-pair mitochondrial cytochrome c oxidase 1 gene (COI) is regarded as the standardized barcode gene for species identification (iBOL 2018). Thus far, there is a mixed record of success of using COI sequences for these purposes for mosquitoes. In an early application of this approach to mosquito identification, Kumar et al. (2007) analyzed 63 species from 15 genera found in India, successfully identifying 62 species. However, they were unable to distinguish between *Ochlerotatus portonovoensis* and *O. wardi*, which are considered closely related species based on morphology (Reinert et al. 2004). Curiously, Kumar et al. (2007) presented a phylogenetic tree of their species, which they claimed (Kumar et al. 2007, pg. 7), “was in general agreement with the taxonomy based on morphology as reported previously”, although it contained several glaring discrepancies from traditional taxonomic schemes. Notably, no *Aedes, Culex*, or *Ochlerotatus* were recovered as monophyletic, yet these issues were not explicitly identified. Chan et al. (2014) analyzed 45 species from 13 genera found in Singapore, and reported a 100% success rate in identifying mosquito species. Similar to Kumar et al. (2007), they presented phylogenetic trees but once again did not draw attention to apparent discrepancies. In the phylogeny composed of *Aedes, Verrallina*, and *Ochlerotatus* species, the sole *Verrallina* representative (*V. butleri)* was clustered together with an *Aedes* species (*A. collessi*) and an *Ochlerotatus* species (*O. cogilli*) in an interior clade, rendering *Aedes* non-monophyletic. Additionally, the two *Ochlerotatus* species included (*O. cogilli, O. vigilax*) did not form a monophyletic grouping. Most recently, Chu et al. (2016) constructed phylogenetic trees for 34 mosquito species using complete mitogenomes, as well as the COI barcoding gene. Although their investigation was not designed to resolve the validity of the genus *Ochlerotatus*, with the majority of sequences representing the genus *Anopheles*, it contained three *Aedes* species and one *Ochlerotatus* species. Relevant to our questions, the genus *Aedes* was not found to be monophyletic relative to *O. vigilax* (or *Haemagogus janthinomys*) in either dataset.

Herein we make use of publicly available DNA barcode data to assess the validity of the genus *Ochlerotatus* relative to *Aedes*, following Reisen’s (2016) call to action. Unlike previous investigations, which overrepresented other taxa (notably *Anopheles* or *Culex*), we specifically targeted *Aedes* and *Ochlerotatus* sequences with the explicit goal of testing their relative monophyly, using an appropriate outgroup (*Anopheles*). We predicted that if *Ochlerotatus* is a valid evolutionary unit, minimally as a subgenus, that our included *Ochlerotatus* species should cluster together separate from *Aedes* with high bootstrap support.

## Methods

### Species Selection

All current and previous genus and species names were confirmed using the literature on Aedini taxonomy (e.g., Wilkerson et al. 2015). The group we call “True *Aedes*” are those species that have previously been classified in the *Aedes* genus and were not part of the *Ochlerotatus* subgenus or reclassified into the new *Ochlerotatus* genus by Reinert et al. (2000, 2004, 2006). The “*Ochlerotatus*” group comprises those species that were previously part of the *Ochlerotatus* subgenus of *Aedes* or were reclassified into the new *Ochlerotatus* genus by Reinert et al. (2000, 2004, 2006). The genus *Anopheles* was selected as the outgroup for the analysis because it is a separate genus that is part of the same family (Culicidae) as *Aedes/Ochlerotatus*.

### Obtaining Sequences

Sequences were downloaded from the Barcode of Life Data (BOLD) Systems (Ratnasingham and Hebert 2013) in FASTA file format and later compiled into one master file, comprising a total of 873 sequences. The sequences cover the COI barcode region of the mitochondrial genome, spanning approximately 650 base pairs. All *Aedes* and *Ochlerotatus* species that had repositories on BOLD were downloaded, but we excluded those that had less than three sequences available for a given species to ensure the sample size was large enough to represent that species. The maximum number of sequences used for each species was 20. In most cases, the first 20 sequences were selected and downloaded. Otherwise, the sequence files were viewed in Molecular Evolutionary Genetic Analysis (MEGA) version 7 (Kumar et al. 2015), and the ones with the most coverage were randomly selected. All of the sequences were manually inspected in MEGA, and those which had unknown “N” bases, missing data, or were not properly aligned were removed from the analysis. In total, 873 sequences were used for analysis (Table 1).

**Table 1.** Total number of species and sequences for each of the three groups used in the analysis of *Aedes-Ochlerotatus* mosquitoes.

|  | **True *Aedes*** | ***Ochlerotatus*** | **Outgroup** |
| --- | --- | --- | --- |
| **Number of Species** | 21 | 36 | 8 |
| **Number of Sequences** | 269 | 501 | 103 |

### Phylogenetic Analyses

Maximum Likelihood (ML), Neighbor Joining (NJ), and Bayesian Inference (BI) methods were employed to generate phylogenies, which were then visualized in FigTree version 1.4.2 (Rambaut 2009). This approach allowed us to compare the congruence of resultant trees. The command line program Randomized Accelerated Maximum Likelihood (RAxML) version 8.0.0 was used to generate the Maximum Likelihood phylogenetic tree (Stamatakis 2014). A bootstrap analysis with 1000 replicates was performed using the sequence master file with all 873 sequences. This tree was inspected to confirm monophyly of species, and those species that failed to be monophyletic were removed from the analysis. Next, one representative sequence from each monophyletic species was selected and the above bootstrap analysis was replicated to generate a ML consensus tree. The trees generated with the master file and with one sequence per species were compared to confirm there were no changes in topology of the tree. Ultimately, 26 True *Aedes*, 37 *Ochlerotatus*, and eight *Anopheles* outgroup species were selected for further analyses.

With the same selected sequences from above, a Bayesian Inference analysis with corresponding posterior probability support values was generated using MrBayes version 3.2.6 (Ronquist et al. 2012) for 1,000,000 generations. Rate heterogeneity was estimated using a gamma distribution model for the variable sites and the first 25% of samples were discarded as burnin. Because only one outgroup could be specified in the program, *Anopheles marajoara* was randomly selected from the *Anopheles* species to be listed as the outgroup. Finally, a Neighbor Joining tree was generated in MEGA version 7 (Kumar ey al. 2015) using a Kimura two-parameter model. Bootstrap values were also calculated with 1,000 replicates.

## Results

Phylogenetic trees were generated via ML, NJ, and BI methods. Using all 873 COI barcode sequences for a ML analysis, we determined that a majority of species clustered together as monophyletic (S1 Fig.). Using one representative sequence from each species, we generated consensus phylogenetic trees (Fig. 1, Fig. 2, Fig. 3). As expected, almost all subspecies were recovered and clustered together with significant support values; *A. aegypti* and *A. aegypti aegypti* (ML 100, BI 100, NJ 100), *A. flavopictus downsi* and *A. flavopictus miyarai* (ML 100, BI 99.2, NJ 100), and *A. vexans* and *A. vexans nipponii* (ML 80, BI 94.8, NJ 100). Conversely, *A. japonicus* and *A. japonicus yaeyamensis* were positioned one node away (ML 96, BI 41.3, NJ 100), clustering *A. japonicus yaeyamensis* with *A. koreicus* with generally lower support values (ML 63, BI 61.1, NJ 88). Finally, we note that even when we did not specify them as constituting an outgroup (i.e., unrooted phylogeny), *Anopheles* was monophyletic and sister to the group containing the remaining sequences.

**Fig 1.**
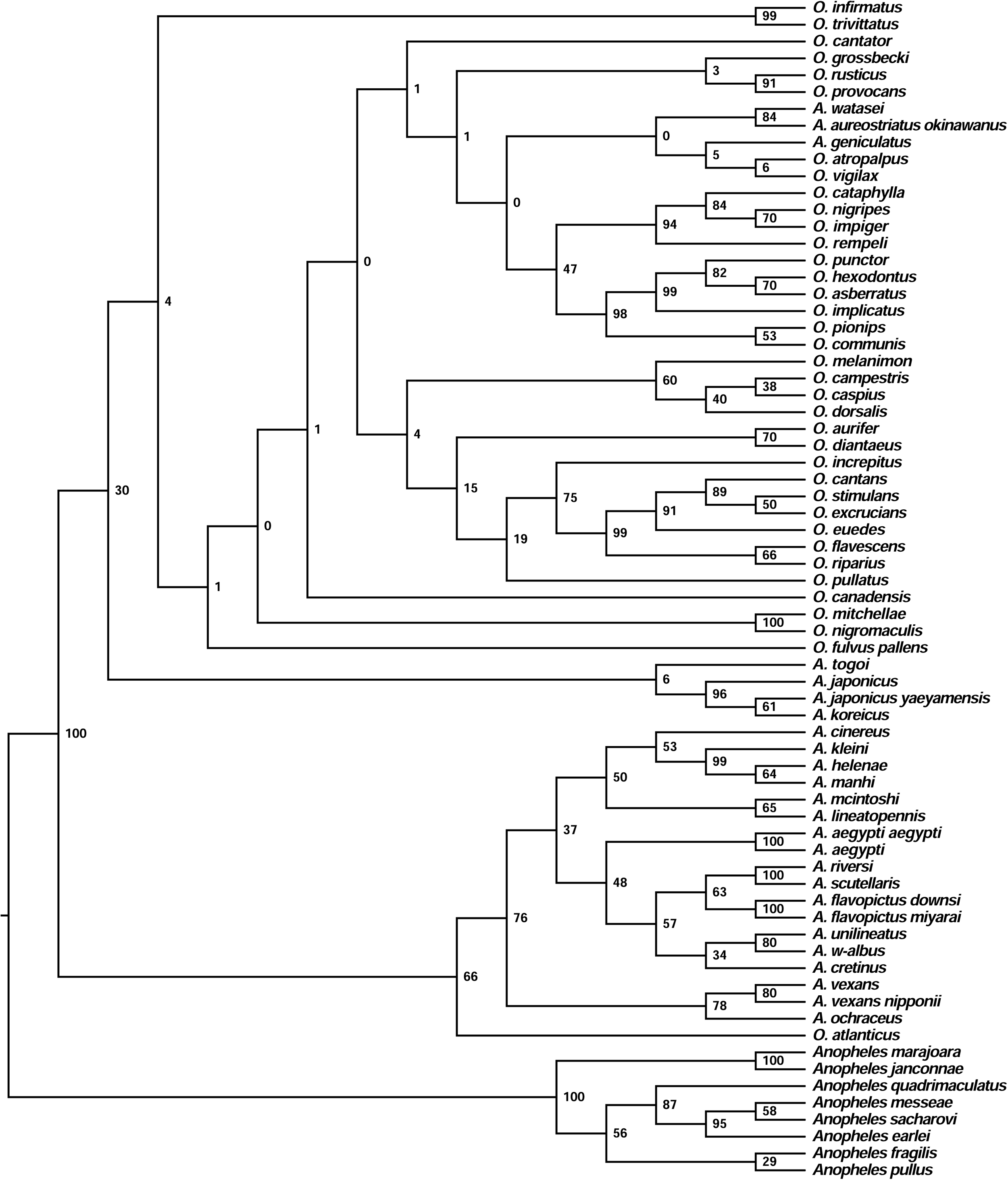
Maximum Likelihood phylogeny of Aedini species based on COI barcoding sequences. Phylogeny generated with RAxML version 8.0.0 (Stamatakis 2014) of True *Aedes, Ochlerotatus*, and outgroup species. A. is *Aedes* genus and O. is *Ochlerotatus* genus. Numbers at each node represent bootstrap values. Tree visualized in FigTree version 1.4.2 (Rambaut 2009).

**Fig 2.**
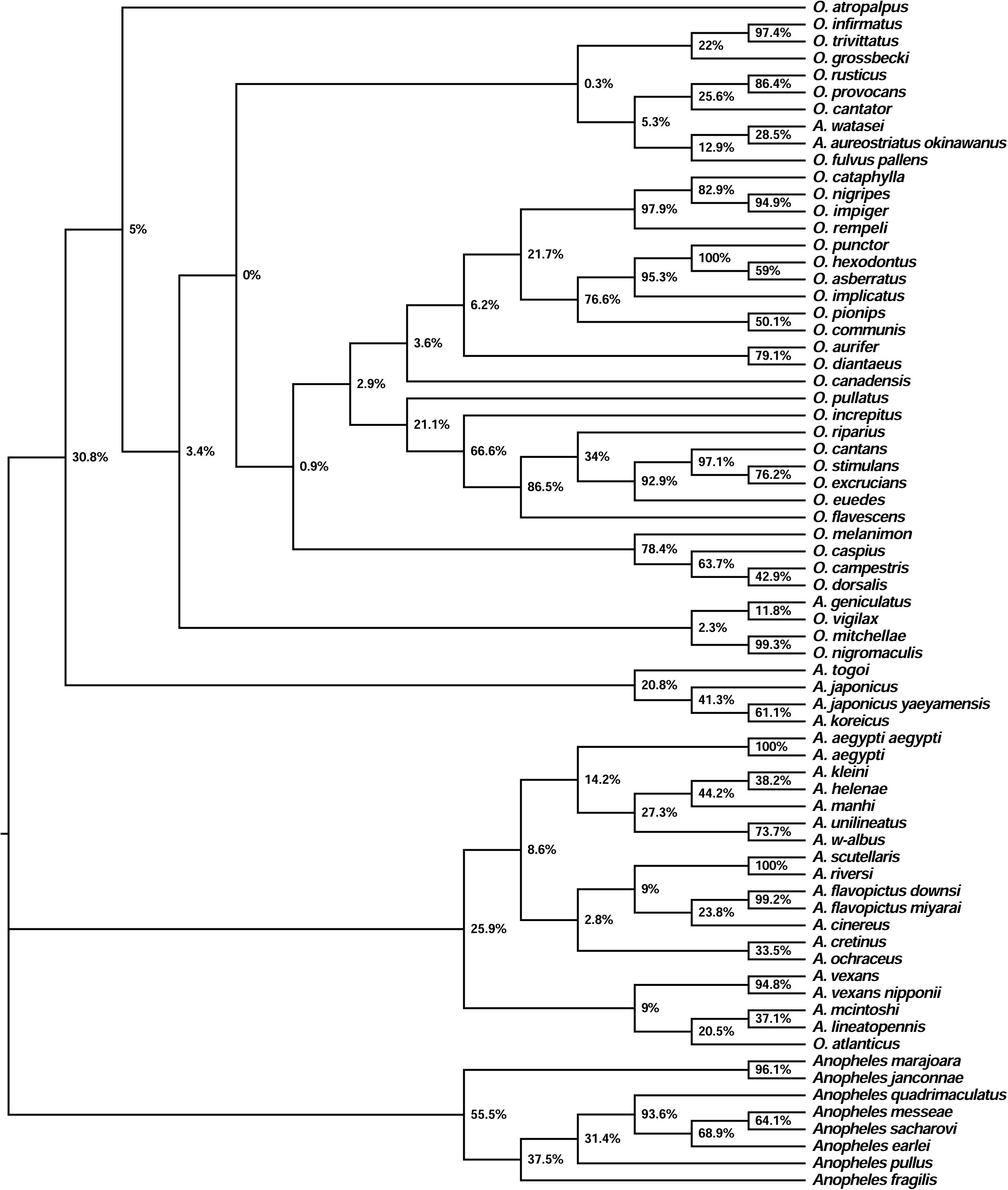
Bayesian Inference phylogeny of Aedini species based on COI barcoding sequences. Phylogeny generated with MrBayes version 3.2.6 (Ronquist et al. 2012) of True *Aedes, Ochlerotatus*, and outgroup species. A. is *Aedes* genus and O. is *Ochlerotatus* genus. Percentages at each node represent posterior probabilities. Tree visualized in FigTree version 1.4.2 (Rambaut 2009).

**Fig 3.**
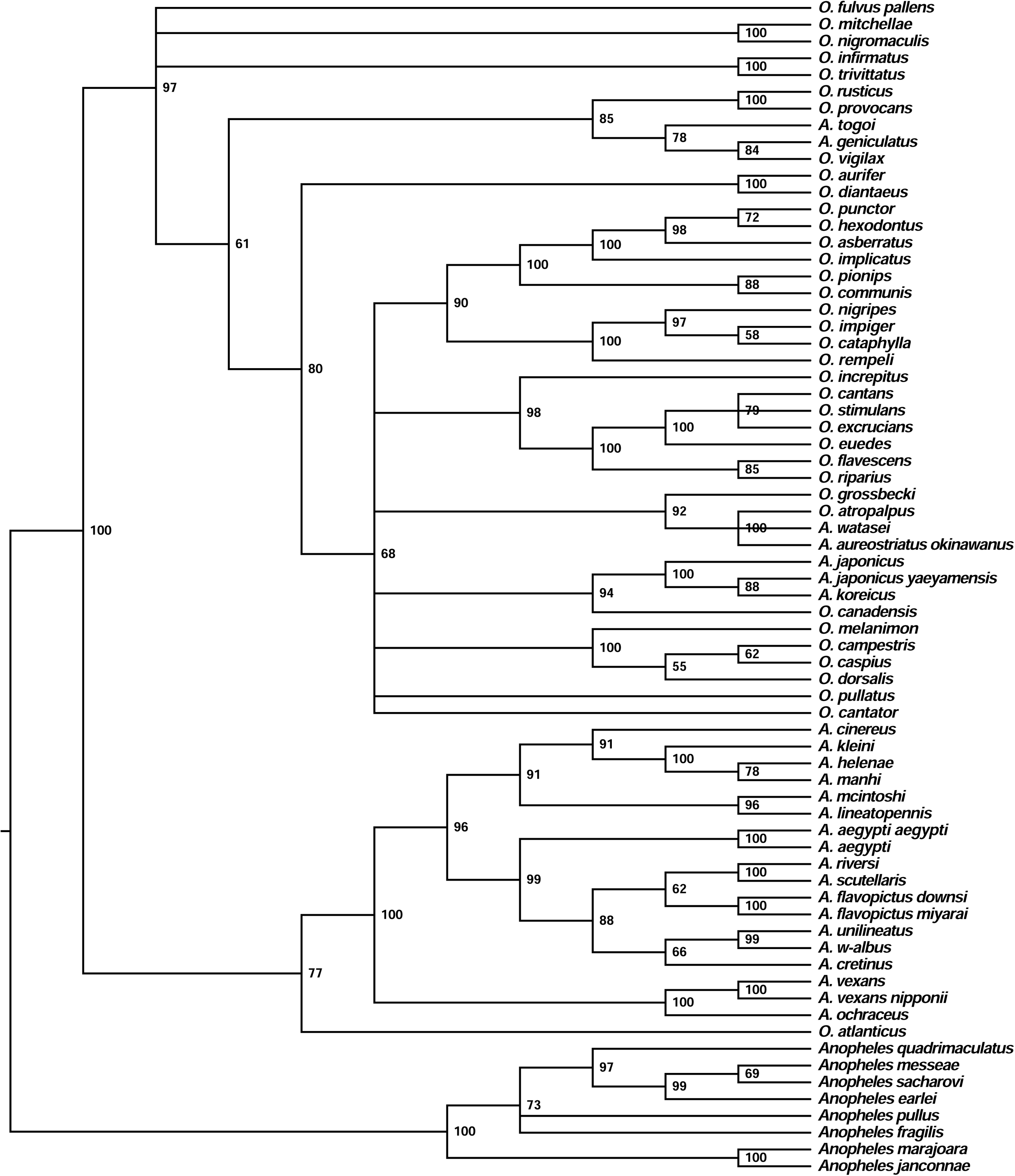
Neighbor Joining phylogeny of Aedini species based on COI barcoding sequences. Phylogeny generated with MEGA version 7 (Kumar et al. 2015) of True *Aedes, Ochlerotatus*, and outgroup species. A. is *Aedes* genus and O. is *Ochlerotatus* genus. Numbers at each node represent bootstrap values. Tree visualized in FigTree version 1.4.2 (Rambaut 2009).

With regard to our main question, the resulting trees did not totally align with the morphology-based classifications previously suggested by Reinert et al. (2000, 2004, 2006, 2008, 2009). Overall, neither *Aedes* nor *Ochlerotatus* were monophyletic in any of the phylogenies generated (Fig. 1, Fig. 2, Fig. 3, S1 Table). We found evidence of non-random clustering consistent across all three phylogenies: (i) in one major group, there was only a single *Ochlerotatus* species (*O. atlanticus*), which was sister to 18 *Aedes* taxa in the ML, BI, and NJ trees, (ii) the remaining *Ochlerotatus* species (N = 36) were contained in the other major group, which were (iii) consistently associated with seven *Aedes* taxa (*A. japonicus, A. japonicus yaeyamensis, A. koreicus, A watasei, A. aureostriatus okinawanus, A. togoi, A. geniculatus*) with generally good agreement between the methods in relative positioning among four of the seven *Aedes* species (Fig. 1, Fig. 2, Fig. 3). By treating the former as the ‘*Aedes* clade’ and the latter as the ‘*Ochlerotatus* clade’, we performed *post hoc* contingency tests to assess the strength of these associations. Because of the high congruence in the tree resulting from the three methods, we used only the ML tree to prevent pseudoreplication. By doing so, we find that the ‘*Aedes* clade’ contained significantly fewer *Ochlerotatus* species than expected by chance (Fisher’s exact test, *P* = 0.003). Similarly, the ‘*Ochlerotatus* clade’ contained significantly fewer *Aedes* species than expected by chance (Fisher’s exact test, *P* < 0.001).

## Discussion

Given the medical importance of mosquitoes within the traditional *Aedes* genus, there is a need for robust data to support revisions to longstanding names for species and to clarify relationships among species. Here, we present our first response to Reisen’s (2016) call to bring genetic data to bear on morphologically-based species groupings. With extensive debate surrounding the genera *Aedes* and *Ochlerotatus*, our analysis attempted to clarify the phylogeny of these groups using molecular data and to compare our results to phylogenies obtained through morphological characteristics. Our analyses produced three main findings in each of the three phylogenetic methods utilized: (1) Sequences associated with a particular species were generally found to cluster together; (2) *Aedes* and *Ochlerotatus* are not reciprocally monophyletic; and (3) Despite the lack of strict monophyly, our *post hoc* analyses support the existence of non-random associations among *Aedes* and *Ochlerotatus* “congeners” in our dataset. The latter finding suggests that some “*Ochlerotatus”* species may form a valid evolutionary unit, although we find insufficient evidence to support its retention as a genus, which echoes conclusions made by Wilkerson et al. (2015) and Soghigian et al. (2017) based on phenotypic and molecular data, respectively.

The observed lack of reciprocal monophyly of *Aedes* and *Ochlerotatus* may result in part, but not completely, from relatively low support values in parts of all three phylogenies. With regard to the ML tree, bootstrap values of 70 and above correlate with ≥ 95% chance that the suggested clade is valid (Hillis and Bull 1993). Thirty nodes have high bootstrap values greater than 70, whereas 38 nodes fall below this threshold (Fig. 1). The node separating the ‘*Aedes* clade’ from the ‘*Ochlerotatus* clade’ did have high support in both the ML and BI trees (support value = 100 in both cases), although the NJ tree contained a three-way polytomy of these clades together with the outgroup. The low support values associated with some nodes could be a result of the species being too closely related to differentiate, or they could indicate that more molecular markers are needed to analyze these species. The results are unlikely to be based on identification errors, given that we observed only three instances where all 10 sequences for a species or subspecies did not cluster together with high bootstrap support. Due to the difficulty of identifying Aedini mosquitoes based on morphological characteristics, inaccurate species naming in BOLD is plausible. Incorrect species naming in online genetic databases has been found in previous studies (e.g., misidentification of spider mite species was detected in COI sequences downloaded from GenBank (Ros and Breeuwer 2007)). With regard to overall bootstrap support, we note that Chu et al. (2016) also reported low values in their phylogeny, including at basal nodes. Our results suggest that morphology alone is not an accurate representation for mosquito systematics, and that molecular differences among the proposed genera point to additional uncertainty in placement of species.

Previous studies have identified significant limitations in the application of COI barcoding to molecular phylogenetics (Dupuis et al. 2012, Hajibabaei et al. 2007, Moritz and Cicero 2004). Although COI barcoding is generally quite effective at differentiating between species, it is not considered to be appropriate for illuminating deeper evolutionary relationships (Hajibabaei et al. 2007). However, Hajibabaei et al. (2007) suggested that COI barcoding can be used to aid in choosing taxa for phylogenetic analysis, as well as providing greater confidence for shallow evolutionary divergence between species. This indicates that although a COI-based phylogeny such as this should not be interpreted as providing resolution toward the deep evolutionary history of these genera, it can be considered a first step towards a more comprehensive molecular phylogeny (see also: Reece et al. 2008, Zhang and Hewitt 1997). Interestingly, support values do not consistently decline with depth in any of our trees. We suggest that next steps in resolving these genera may be to (1) apply multiple markers, and (2) perform phylogenetic analyses with those genetic data that include phenotypic data as a data partition. Using multiple markers, such as additional nuclear markers commonly used in insect systematics (e.g., *wng*, H3, 18S), would increase the probability of successful delimitation between closely related species, and ultimately generate a more detailed and robust phylogeny (Dupuis et al. 2012). With regard to phenotypic data, there are 336 characters, of which 14 are ordered characters (Wilkerson et al. 2015) that could be represented as a data partition in a Bayesian phylogenetic approach (Drummond et al. 2012). We suggest that an analysis with a greater number of genetic markers, possibly including phenotypic data, be performed for these species for a more accurate representation of their phylogenetic relationships.

Subsequent to commencing our study, we became aware of a related investigation by Soghigian et al. (2017). Soghigian et al. (2017) were particularly interested in the spread of invasive *Aedes* mosquitoes, and attempted to reconstruct the evolution of habitat specialization within the larger group (Aedini). Accordingly, they did not set out to resolve the separation of *Aedes* from *Ochlerotatus*, both of which were treated as *Aedes* subgenera, but their findings are relevant to our overall conclusions. They recovered two major clades with high levels of support, with *Ochlerotatus* being part of Clade B and *Aedes* part of Clade A. However, similar to our findings, *Ochlerotatus* was not monophyletic within its clade. Furthermore, Clade A contained other aedine genera, rendering the genus *Aedes* itself non-monophyletic. Collectively, our results (see also Kumar et al. 2007, Chan et al. 2014, Chu et al. 2016) find little evidence for *Ochlerotatus* being a valid genus, or even subgenus as currently described.

Because Aedini mosquitos are vectors for disease such as yellow fever, malaria, dengue, and West Nile, having confidence in their phylogenetic relationships has implications for public health management. DNA barcoding is an easy standardized method that is an inexpensive way to account for genetics in taxonomy. In this case, there is little need for specialists to make morphology-based species identifications, because species identities can be based on DNA at any life stage. A genetic barcoding approach should serve as an additional tool for taxonomists to supplement their knowledge as well as being an innovative device for non-experts who need to make a quick identification. Ultimately, entomologists should incorporate both morphology and genetics into species classification analyses, which has never been done before.

## Acknowledgements

Author contributions: Methodology, Formal Analysis, Visualization, Project Administration: HG. Data Curation, Resources: EC, DC, HK. Conceptualization, Supervision: SMV, DY. Writing-Original Draft Preparation: HG, EC, DC, HK, SMV. Writing-Review & Editing: HG, SMV, DY.

## Figure and Supporting Information Captions

**S1 Fig. Maximum Likliehood phylogenetic tree of all 873 sequences.** ML phylogeny generated by RAxML version 8.0.0 as a bootstrap analysis with 1000 replicates and visualized in FigTree (Stamatakis 2014, Rambaut 2009). Each branch tip states the accession number and default species names from BOLD and bootstrap values are at each node. Note that many species are named *Aedes* here because they were uploaded to BOLD before those species were reclassified to *Ochlerotatus*. Genus names in Fig 1 (in text) were changed to reflect species name reclassification.

**S1 Table. Species name and accession number for the sequences used in the analysis.** One sequence from each species was randomly selected from the 873 total sequences, and these sequences were used in the bootstrap analysis to generate Fig 1.

