## Supplementary figures and images for "A barcoding approach to phylogenetic classification of Aedini mosquitoes (*Aedes, Ochlerotatus*)"

### Supplementary Materials

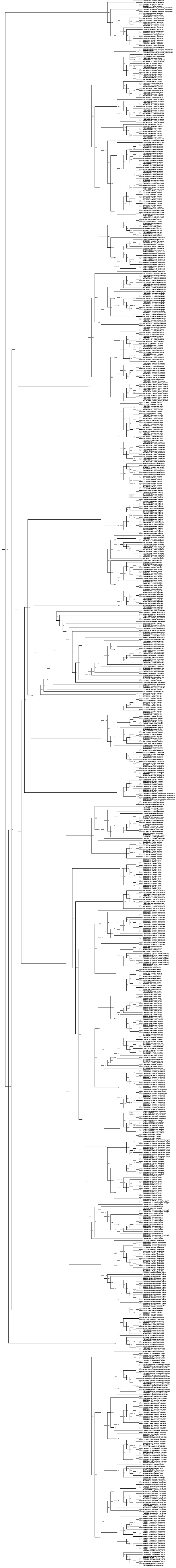
