## Supplementary Materials for "A barcoding approach to phylogenetic classification of Aedini mosquitoes (*Aedes, Ochlerotatus*)"

| Species name | BOLD Sequence Identifier |
|---|---|
| *Aedes aegypti* | GBDCU1629-15\|Aedes_aegypti\|COI-5P\|KM613073 |
| *Aedes aegypti aegypti* | GBDCU1587-15\|Aedes_aegypti_aegypti\|COI-5P\|KF564652 |
| *Aedes aureostriatus okinawanus* | GBDP13510-13\|Aedes_aureostriatus_okinawanus\|COI-5P\|AB738128 |
| *Aedes cinereus* | CNKOI227-14\|Aedes_cinereus\|COI-5P\|KR382490 |
| *Aedes cretinus* | MBIG147-09\|Aedes_cretinus\|COI-5P\|KC250446 |
| *Aedes flavopictus downsi* | GBDP13502-13\|Aedes_flavopictus_downsi\|COI-5P\|AB738136 |
| *Aedes flavopictus miyarai* | GBDP13397-13\|Aedes_flavopictus_miyarai\|COI-5P\|AB738213 |
| *Aedes geniculatus* | CULBE077-14\|Aedes_geniculatus\|COI-5P\|KM258314 |
| *Aedes helenae* | GBDCU397-12\|Aedes_helenae\|COI-5P\|HQ398919 |
| *Aedes japonicus* | GBDCU189-12\|Aedes_japonicus\|COI-5P\|FJ641869 |
| *Aedes japonicus yaeyamensis* | GBDP13400-13\|Aedes_japonicus_yaeyamensis\|COI-5P\|AB738210 |
| *Aedes kleini* | GBDCU391-12\|Aedes_kleini\|COI-5P\|HQ398913 |
| *Aedes koreicus* | CULBE093-14\|Aedes_koreicus\|COI-5P\|KM258298 |
| *Aedes lineatopennis* | GBDCU387-12\|Aedes_lineatopennis\|COI-5P\|HQ398909 |
| *Aedes manhi* | GBDCU402-12\|Aedes_manhi\|COI-5P\|HQ398924 |
| *Aedes mcintoshi* | GBDCU1723-15\|Aedes_mcintoshi\|COI-5P\|KJ940563 |
| *Aedes ochraceus* | GBDCU1870-15\|Aedes_ochraceus\|COI-5P\|KJ940705 |
| *Aedes riversi* | GBDP13357-13\|Aedes_riversi\|COI-5P\|AB738253 |
| *Aedes scutellaris* | GBDP16997-15\|Aedes_scutellaris\|COI-5P\|KP843376 |
| *Aedes unilineatus* | MAMOS087-12\|Aedes_unilineatus\|COI-5P\|KF406626 |
| *Aedes vevans niponii* | GBDP13539-13\|Aedes_vexans_nipponii\|COI-5P\|AB738099 |
| *Aedes vexans* | ACMC053-04\|Aedes_vexans\|COI-5P\|GU907990 |
| *Aedes w-albus* | MAMOS132-12\|Aedes_w-albus\|COI-5P\|KF406649 |
| *Aedes watasei* | GBDP13291-13\|Aedes_watasei\|COI-5P\|AB738138 |
| *Anopheles earlei* | BBGCO1108-15\|Anopheles_earlei |
| *Anopheles fragilis* | GBDP17132-15\|Anopheles_fragilis\|KF564689 |
| *Anopheles janconnae* | MBII348-09\|Anopheles_janconnae\|JQ615430 |
| *Anopheles marajoara* | ALBIS015-12\|Anopheles_marajoara\|KJ011976 |
| *Anopheles messeae* | GBDP11016-12\|Anopheles_messeae\|HE659591 |
| *Anopheles pullus* | GBDP14474-13\|Anopheles_pullus\|KC855646 |
| *Anopheles quadrimaculatus* | IUP957-14\|Anopheles_quadrimaculatus |
| *Anopheles sacharovi* | GBDP2232-06\|Anopheles_sacharovi\|DQ118170 |
| *Ochlerotatus abserratus* | JWDCC270-10\|Aedes_abserratus\|COI-5P\|HM861483 |
| *Ochlerotatus atlanticus* | NEONU1534-12\|Aedes_atlanticus\|COI-5P |
| *Ochlerotatus atropalpus* | BBDCP589-10\|Aedes_atropalpus\|COI-5P\|JF868951 |
| *Ochlerotatus aurifer* | NEONW109-11\|Aedes_aurifer\|COI-5P\|JX259520 |
| *Ochlerotatus campestris* | SAMOS008-09\|Aedes_campestris\|COI-5P\|KR749416 |
| *Ochlerotatus canadensis* | ACMC184-04\|Aedes_canadensis\|COI-5P\|GU907859 |
| *Ochlerotatus cantans* | CULBE019-14\|Aedes_cantans\|COI-5P\|KM258371 |
| *Ochlerotatus cantator* | BBDEE797-10\|Aedes_cantator\|COI-5P\|KR437907 |
| *Ochlerotatus caspius* | CULBE029-14\|Aedes_caspius\|COI-5P\|KM258356 |
| *Ochlerotatus cataphylla* | NEONU348-11\|Aedes_cataphylla\|COI-5P |

| | |
|---|---|
| *Ochlerotatus communis* | ACMC038-04\|Aedes_communis\|COI-5P\|GU907873 |
| *Ochlerotatus diantaeus* | NEONV131-11\|Aedes_diantaeus\|COI-5P\|JX259584 |
| *Ochlerotatus dorsalis* | BBDCP592-10\|Aedes_dorsalis\|COI-5P\|JF868954 |
| *Ochlerotatus euedes* | ACMC126-04\|Aedes_euedes\|COI-5P\|GU907883 |
| *Ochlerotatus excruians* | >JWDCA320-10\|Aedes_excrucians\|COI-5P\|HQ569473 |
| *Ochlerotatus flavescens* | JWDCF258-10\|Aedes_flavescens\|COI-5P\|JF875088 |
| *Ochlerotatus fulvus pallens* | NEONU1609-12\|Aedes_fulvus_pallens\|COI-5P |
| *Ochlerotatus grossbecki* | ACMC022-04\|Aedes_grossbecki\|COI-5P\|GU907901 |
| *Ochlerotatus hexodontus* | JWDCJ822-11\|Aedes_hexodontus\|COI-5P\|KR665263 |
| *Ochlerotatus impiger* | JWDCJ447-11\|Aedes_impiger\|COI-5P\|JF878024 |
| *Ochlerotatus implicatus* | ACMC090-04\|Aedes_implicatus\|COI-5P\|GU907904 |
| *Ochlerotatus increpitus* | NEONU366-11\|Aedes_increpitus\|COI-5P |
| *Ochlerotatus infirmatus* | NEONU1563-12\|Aedes_infirmatus\|COI-5P |
| *Ochlerotatus melanimon* | NEONU461-11\|Aedes_melanimon\|COI-5P |
| *Ochlerotatus mitchellae* | NEONU1541-12\|Aedes_mitchellae\|COI-5P |
| *Ochlerotatus nigripes* | GMGLC1066-13\|Aedes_nigripes\|COI-5P |
| *Ochlerotatus nigromaculis* | NEONU298-11\|Aedes_nigromaculis\|COI-5P |
| *Ochlerotatus pionips* | MMADL011-09\|Aedes_pionips\|COI-5P\|GU680395 |
| *Ochlerotatus provocans* | CNGBK137-14\|Aedes_provocans\|COI-5P\|KR383704 |
| *Ochlerotatus pullatus* | NEONU334-11\|Aedes_pullatus\|COI-5P |
| *Ochlerotatus punctor* | CULBE109-14\|Aedes_punctor\|COI-5P\|KM258293 |
| *Ochlerotatus rempeli* | SAMOS038-09\|Aedes_rempeli\|COI-5P\|KR758245 |
| *Ochlerotatus riparius* | JWDCK160-11\|Aedes_riparius\|COI-5P\|KR655256 |
| *Ochlerotatus rusticus* | CULBE120-14\|Aedes_rusticus\|COI-5P\|KM258273 |
| *Ochlerotatus stimulans* | ACMC031-04\|Aedes_stimulans\|COI-5P\|GU907950 |
| *Ochlerotatus trivittatus* | ACMC103-04\|Aedes_trivittatus\|COI-5P\|GU907982 |
| *Ochlerotatus vigilax* | GBDCU447-12\|Ochlerotatus_vigilax\|COI-5P\|JN228485 |
